# Dual transcriptomics reveal commensal interactions between microalgae and phycosphere bacteria

**DOI:** 10.64898/2026.02.09.704904

**Authors:** Line Roager, Morten Dencker Schostag, Alessandro N. Garritano, Lone Gram, Suhelen Egan

## Abstract

The interactions between microalgae and the bacteria living in the phycosphere are pivotal to the role they play in aquatic ecosystems. This study examines how two representatives of common phycosphere bacteria, *Yoonia* sp. TsM2_T14_4 (Rhodobacteraceae) and *Maribacter* sp. IgM3_T14_3 (Flavobacteriaceae), interact with three microalgal hosts: *Isochrysis galbana*, *Tetraselmis suecica*, and *Conticribra weissflogii* (formerly *Thalassiosira weissflogii*) using dual transcriptomic analyses of both bacteria and microalgae. Bacterial transcriptomes differed significantly depending on microalgal host, with notable changes in carbohydrate metabolism among other COG categories. *Yoonia* sp. expressed genes involved in anoxygenic photosynthesis in co-culture with *I. galbana*, presumably due to its inability to utilize carbohydrates from this algal host, whereas *Maribacter* sp. expressed polysaccharide degradation genes in co-culture with *C. weissflogii* along with T9SS genes, which can be employed to secrete these hydrolytic enzymes. Specifically, a putative glucan endo-1,3-beta-D-glucosidase was highly expressed, an enzyme that can hydrolyze laminarin and curdlan. *Maribacter* sp. IgM3_T14_3 could utilize laminarin as a sole carbon source in laboratory settings, a polysaccharide commonly found in marine environments and produced by *C. weissflogii*. Surprisingly, microalgal transcriptomes remained largely unaltered in the presence of either of the bacteria compared to transcriptomes of axenic algal cultures. These findings highlight the adaptability of phycosphere bacteria to different microalgal hosts. Furthermore, it also indicates a commensalism between microalgae, *Yoonia* sp. and *Maribacter* sp., in which the bacteria adapt to and benefit from microalgal host exudates, whereas under the conditions employed here the microalgae are unaffected by the presence of these bacterial symbionts.

**Importance:** Microalgae are the key players in marine ecosystems, capturing carbon dioxide through photosynthesis and releasing carbohydrates into their immediate environment, the so-called phycosphere. Certain bacterial taxa are consistently found within the phycosphere, where they interact with their microalgal host in a variety of ways. However, the impact of these bacteria on the microalgae is not fully understood despite their ecological relevance. This study uses a dual transcriptomic approach to investigate the impact of such core phycosphere bacteria on microalgal hosts and vice versa to uncover the reason behind their success in the phycosphere and possible roles in marine ecosystems.

## Introduction

Microalgae are the main primary producers of the oceans and, thus, cornerstones in oceanic ecosystems, where they interact with zooplankton, bacteria, and viruses. Microalgal-bacterial interactions in the phycosphere, the microenvironments surrounding microalgae (1, 2), have received increasing attention in recent years as major drivers of biogeochemical cycles as well as for the use of algae in biotechnology, aquaculture, and biofuel production (3–6).

The interactions between algae and bacteria are complex, involving a range of mutualistic, commensal, and sometimes parasitic relationships, often with multiple partners (7–16). For instance, a *Bacillus* endophyte of *Ulva* sp. enhances the presence of other phycosphere bacteria, which in turn protects against the pathogen *Aquimarina* (17). Other bacteria switch from mutualistic or commensal lifestyles to pathogenic ones as their algal hosts enter senescence (18, 19). This is attributed to the release of certain algal metabolites and dissolved organic matter (DOM), which generally shape the composition of the phycosphere microbiome (20–22), leading to the proposal that microalgal hosts select for beneficial microbiomes (23).

In a previous study (24), we observed host-specific phycosphere microbiome communities, forming from the same original seawater microbiome, in microalgal cultures of *Isochrysis galbana, Tetraselmis suecica,* and *Conticribra weissflogii*. Microbiomes were dominated by Rhodobacteraceae and Flavobacteriaceae, with Rhodobacteraceae especially prevalent in *T. suecica* and *C. weissflogii* microbiomes, and Flavobacteriaceae dominating *I. galbana* microbiomes. Other studies have reported similar findings, where Rhodobacteraceae and Flavobacteriaceae have repeatedly been found in association with microalgae, both in laboratory settings and natural blooms (23, 25–34). Flavobacteriaceae are capable of degrading high-molecular weight (HMW) carbohydrates and have been proposed as degraders of microalgal DOM in the decline phase of phytoplankton blooms, and in some cases proposed as algicidal agents causing the end of a bloom (28, 32, 35, 36). Rhodobacteraceae are often found in association with algae (both micro- and macroalgae) and can peak in abundance in the beginning of a phytoplankton bloom (25, 27–30). In contrast to Flavobacteriaceae, Rhodobacteraceae have poor polysaccharide degradation capabilities, and generally consume low-molecular weight (LMW) carbohydrates (37–40). It is currently unknown whether members of these bacterial families are so frequently abundant in phycospheres because they confer benefits to the microalgal hosts. If so, this would be an incentive for the microalgae to actively select for these bacteria in their phycosphere microbiomes.

The purpose of the present study was to investigate if phycosphere bacteria from these taxa confer benefits to microalgal hosts and thus might be selected for. Furthermore, to investigate which benefits the phycosphere bacteria may receive from the microalgal hosts. For this purpose, we selected two representatives of Flavobacteriaceae and Rhodobacteraceae isolated from our previous study (24); a *Maribacter* sp. and a *Yoonia* sp., and the microalgae *I. galbana, T. suecica,* and *C. weissflogii*. These bacteria were members of the core communities of their respective hosts, i.e. *Maribacter* sp. with *I. galbana,* and *Yoonia* sp. with *T. suecica* and *C. weissflogii.* In contrast to other studies which have investigated the transcriptional response of either phycosphere bacteria (11, 31, 41–44) or microalgal hosts (31), we investigate both the gene expression profiles of the bacteria and the microalgal hosts, aiming to determine if and how the microalgal host benefits from the presence of these core phycosphere bacteria and vice versa. Benefits seen in the microalgal transcriptome profile could be in the form of lessened stress signals, increased photosynthetic productivity, transport of nutrients, or Calvin cycle enzyme activity. Similarly, benefits to the phycosphere bacteria may be reflected in the transcriptome in terms of decreased stress signals, increased uptake of nutrients or carbohydrates etc. With this analysis, we aim to gain insight into the reasons for the success of these representatives of Rhodobacteraceae and Flavobacteriaceae in the phycosphere. Are these bacteria conferring benefits to their microalgal hosts and vice versa, or is the interaction beneficial to one of the parties only?

## Materials and Methods

### Bacterial and microalgal strains and culture

*Yoonia* sp. TsM2_T14_4 and *Maribacter* sp. IgM3_T14_3 were isolated on marine agar (MA, Difco 2216) plates from microbiomes associated with the microalgal hosts *Tetraselmis suecica* CCAP 66/4 and *Isochrysis galbana* CCMP1323. The host-specific microbiomes of these two algae originated from the same natural seawater as previously described (45). Both bacterial isolates were routinely grown on MA or in marine broth (MB, Difco) at 25 °C and 200 rpm. Axenic cultures of *I. galbana* CCMP1323 and *T. suecica* CCAP 66/4 were grown in f/2 medium (46, 47) without silica, based on a 3% Instant Ocean (IO, Aquarium Systems) solution, which will be referred to as f/2-Si 3% IO. Axenic *Conticribra weissflogii* CCMP1336 was grown in f/2+Si 3% IO. All microalgae were cultivated at 18 °C and constant light at ∼50 µmol m^-2^ s^-1^.

### Bacterial genome sequencing, assembly, and annotation

*Yoonia* sp. TsM2_T14_4 and *Maribacter* sp. IgM3_T14_3 were grown for 2 days as described above before DNA extraction. Briefly, cultures were pelleted at 8,000 × g for 5 min before being resuspended in a sucrose lysis buffer (400 mM NaCl, 750 mM sucrose, 20 mM EDTA, 50 mM Tris-HCl). Lysozyme was added (final conc. 10 mg/ml), and samples were incubated at 37 °C for 30 min. Then, proteinase K (final conc. 100 µg/ml) and SDS (final conc. 1% w/v) were added before incubation overnight (O/N) at 55 °C and 300 rpm. Lysed samples were added to Maxwell^®^ RSC Cartridges before automatic DNA extraction in a Maxwell^®^ Instrument (Promega) using the Maxwell^®^ RSC Blood DNA Kit. Extracted DNA was eluted in the kit elution buffer before measurement of DNA quantity and concentration using a Qubit 2.0 Fluorometer and the broad range (BR) assay kit (Thermo Fisher Scientific). DNA quality was also assessed using a Denovix DS-11 spectrophotometer. DNA was sequenced (Novogene, Cambridge, UK) on an Illumina NovaSeq 6000 instrument with 2 x 150bp chemistry. In addition, long-read sequencing was performed in-house on a MinION device (Oxford Nanopore Technologies) using a Rapid Barcoding Sequencing Kit (SQK-RBK004) and a MinION Flow Cell (R9.4.1) according to the appertaining protocols. The resulting raw sequencing data was quality-filtered, and adapters were removed using fastp (v0.23.2) (48, 49) for short reads and porechop (v0.2.4) (50) for long reads. Subsequently, hybrid assembly of short and long reads was performed using Unicycler (v0.5.0) (51) and resulting assemblies were visualized using Bandage (52) and putatively identified using average nucleotide identity by autoMLST (53). Finally, assemblies were annotated using Bakta (v.1.8.2) (54), employing the tools tRNAscan-SE 2.0 (55), Aragorn (56), Infernal (57), PilerCR (58), Prodigal (59), Diamond (60), BLAST+ (61), HMMER (62), and AMRFinderPlus (63), and the databases Rfam (64), Mob-suite (65), DoriC (66), AntiFam (67), UniProt (68), RefSeq (69), COG (70), KEGG (71), PHROG (72), AMRFinder (73), ISFinder (74), Pfam (75), and VFDB (76).

The *Yoonia* sp. genome assembly was mined for genes belonging to photosynthetic gene clusters (PGCs) encoding the apparatus necessary for anoxygenic photosynthesis. PGC amino acid sequences were obtained from a publicly available *Yoonia vestfoldensis* SKA53 genome (GenBank accession no. GCA_000152785.1) and were queried against the assembled *Yoonia* sp. genome of this study using BLASTP with standard settings (61, 77). 18 genes that are known components of PGCs were selected, including *bchBCFLNXYZ, pufABLMQ, ppsR*, and *crtACFI* (78, 79). PGCs found in the *Yoonia* sp. TsM2_T14_2 genome were visualized using the tidyverse and geneviewer packages in R (v4.3.1) (80, 81).

### Microalgal-bacterial co-culture experiment and sampling

A co-culture experiment was set up with the three microalgal hosts *I. galbana, T. suecica,* and *C. weissflogii* and the two bacterial isolates *Yoonia* sp. TsM2_T14_4 and *Maribacter* sp. IgM3_T14_3 (Figure 1). Microalgal pre-cultures were cultivated as described above by diluting outgrown cultures at 10% v/v in fresh media and growing for 7 days before the start of the experiment. Algal cell density was determined using a Neubauer-improved counting chamber, and density was adjusted to ∼10^5^ cells/ml in the final 110 ml co-cultures with fresh f/2±Si 3% IO. The two bacterial isolates were grown as described above for two days prior to the start of the experiment. Thereafter, the cultures were centrifuged for 3 min at 10,000 × g, supernatant removed, the pellet resuspended in 3% IO solution and OD_600_ was adjusted to 0.1 before adding to the co-cultures at 1% v/v. All combinations of host microalgae and bacterial isolate were prepared along with axenic microalgal cultures. All cultures were prepared in biological quadruplicate and incubated at 18 °C with light ∼50 µmol photon m^-2^ s^-1^ until the host microalgae reached exponential growth phase (6 days for *I. galbana* and *T. suecica,* 4 days for *C. weissflogii*). At this point, 1 ml samples were taken from all cultures for quantification of microalgal and bacterial cell densities. Bacterial CFU counts were determined from 10-fold dilution series plated on MA, and the remainder of the undiluted samples were fixed in 1% formaldehyde and stored at 4 °C until algal cell quantification. Algal cell density was determined by flow cytometry (MACSQuant^®^ VYB instrument). In addition, 2 samples of 45 ml from each culture were mixed with ice-cold 5 ml RNA stop solution (5% phenol in absolute ethanol) (82). After centrifugation at 12,000 × g for 5 min, supernatant was discarded (apart from 1 ml) and the two samples from each culture merged and resuspended in 1 ml supernatant. Resuspended samples were then transferred to 1.5 ml bead beat tubes and centrifuged again at 12,000 × g for 5 min. Supernatant was discarded, and pellets were snap-frozen in liquid nitrogen before storage at -80 °C until RNA extraction.

**Figure 1.**
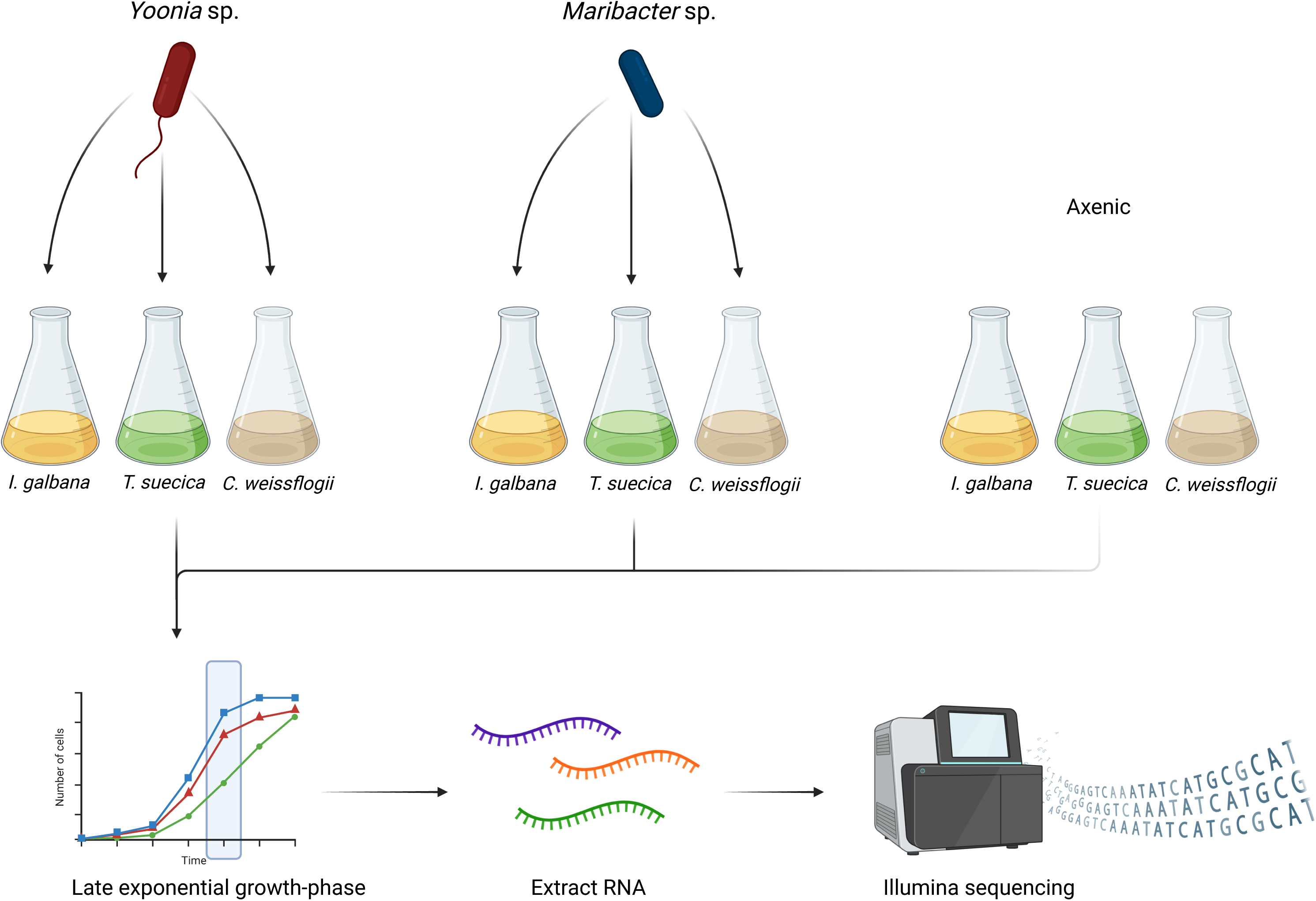
Overview of experimental setup of microalgal-bacterial co-cultures and subsequent transcriptome sequencing. Cultures of the three host microalgae *I. galbana, T. suecica,* and *C. weissflogii* were grown in quadruplicates either axenically or in co-culture with in quadruplicates either axenically or in co-culture with *Yoonia* sp. TsM2_T14_4 or *Maribacter* sp. IgM3_T14_3 until late exponential phase before extraction of RNA and subsequent sequencing. Figure created with BioRender.com.

### RNA extraction and sequencing

RNA was extracted from samples from the co-culture experiments using the RNeasy^®^ PowerPlant^®^ Kit (Qiagen). The protocol from the kit was followed, including the optional addition of 50 µl phenolic separation solution and homogenization using a FastPrep^®^ machine at speed 5.5 for 30 s. Subsequently, residual DNA was removed from samples using the TURBO DNA-*free^™^* Kit (Invitrogen*^™^*). Quality and quantity of extracted RNA was assessed using the RNA high sensitivity assay kit and a Qubit Fluorometer, and the RNA 6000 Nano Kit and a 2100 Bioanalyzer (Agilent). Samples were then shipped to the sequencing provider (Novogene, Cambridge, UK) who depleted samples of rRNA, fragmented RNA, and reverse transcribed into cDNA prior to library preparation and strand-specific sequencing on an Illumina Novaseq 6000 instrument with 2 x 150bp chemistry.

### Data handling and analysis

RNA sequencing data was quality controlled using FastQC (83) before trimming based on quality with trimmomatic (84) along with removing adapter sequences. Subsequently, rRNA sequences were filtered out using SortMeRNA (85), and kallisto (86) was used for pseudoalignment and quantification of reads. Pseudoalignment was done based on the bacterial genome assemblies generated, and for microalgal hosts, in-house available genome assemblies were used. Predicted genes based on genome assemblies were utilized at a protein level for alignment. Quantified reads were imported to R (v4.3.1) using the tximport package (87), and differential expression analysis was performed using DESeq2 (88), and NMDS and PERMANOVA analyses were performed using vegan (89). Results with adjusted p-values of < 0.01 and absolute log2-fold changes of > 1 were further examined and visualized using the tidyverse, ggplot2, and pheatmap packages (80, 90, 91). In addition, an analysis of functional categories was carried out using eggnog-mapper (92, 93), and results were visualized using the aforementioned packages along with ggbreak (94).

### Chitin degradation assay

The possible chitin degradation of *Yoonia* sp. TsM2_T14_2 was tested in a plate-based assay previously employed (95, 96). In brief, 20 µl O/N culture of *Yoonia* sp. TsM2_T14_2 was spotted on agar plates containing 2% IO, 1.5% agar, 0.3% casamino acids, and 0.2% chitin. Both crystalline chitin from crab and colloidal chitin prepared as previously published (96) was tested. A known chitin degrader, *Pseudoalteromonas* sp. S201 (97), was included as a positive control, and clearing zones were inspected after 7 days of growth at 25 °C.

### Cobalamin assay

The possible cobalamin supply of *Yoonia* sp. TsM2_T14_2 to *C. weissflogii* was tested by preparing f/2+Si 3% IO medium with a vitamin solution without cobalamin (f/2+Si - B12 3% IO). A co-culture was set up in this medium with *C. weissflogii* diluted to an approximate starting concentration of ∼10^4^ cells/ml and *Yoonia* sp. TsM2_T14_2 was grown O/N, centrifuged for 3 min at 10,000 × g, supernatant removed, the pellet resuspended in 3% IO and OD_600_ was adjusted to 0.1 before adding at 1% v/v. A control culture with no addition of *Yoonia* sp. TsM2_T14_2 was also set up, as well as a culture in normal f/2+Si 3% IO with all vitamins supplied. Cultures were in biological quadruplicates and were incubated for 20 days at 18 °C at constant light ∼50 µmol m^-2^ s^-1^. Algal growth was followed by flow cytometry, and bacterial density was monitored through CFU counts on MA plates on days 0, 2, 5, 12, and 20.

### Polysaccharide utilization

The ability of *Maribacter* sp. IgM3_T14_3 to utilize alginate, curdlan, and laminarin as sole carbon sources was tested as follows: *Maribacter* was grown in MB for 5 days at 25 °C and 200 rpm before centrifugation at 10,000 × g for 3 min. The supernatant was discarded and the pellet washed in 3% IO twice before resuspension in 3% IO. OD_600_ was measured and *Maribacter* IgM3_T14_3 inoculated at an approximate OD of 0.01 in f/2-Si 3% IO with 1 mM added alginate, curdlan, laminarin, or glucose. Cultures were then incubated at 25 °C and 200 rpm. Utilization of the carbon sources was determined as visible growth observed after 7 and 14 days.

### Data availability

The assembled bacterial genomes are available in GenBank under accession numbers CP187138 (*Maribacter* sp. IgM3_T14_3) and CP187137 (*Yoonia* sp. TsM2_T14_2). Raw RNA sequencing data from co-culture experiments is available in the SRA database under BioProject PRJNA1246571. Finally, scripts for transcriptome analyses are available on github.com/lineroager/phycosphere_dual_transcriptomics.

## Results

### Bacterial genome assemblies

Genomes of *Yoonia* sp. TsM2_T14_4 and *Maribacter* sp. IgM3_T14_3 were assembled and fully closed (1 contig each, all information in Table 1). The IgM3_T14_3 genome is 4,897,888 bp, has a GC% of 35.43, and is most closely related to a *Maribacter aquivivus*, with an ANI of 89.2%. The TsM2_T14_4 genome is 2,960,374 bp, has a GC% of 60.24, and is most closely related to *Yoonia vestfoldensis* with an ANI of 98.8%. Hence, isolate TsM2_T14_4 will be referred to as *Yoonia* sp. TsM2_T14_4, and isolate IgM3_T14_3 will be referred to as *Maribacter* sp. IgM3_T14_3. Based on the annotation of the genomes, *Yoonia* sp. TsM2_T14_4 contained 2891 CDSs, 45 tRNA genes, and 6 rRNA genes. 42 genes were not identified and were annotated as hypothetical proteins. For *Maribacter* sp. IgM3_T14_3, the genome contained 4366 CDSs, 42 tRNA genes, 9 rRNA genes, and 355 genes were unidentified and annotated as hypothetical proteins.

**Table 1.**
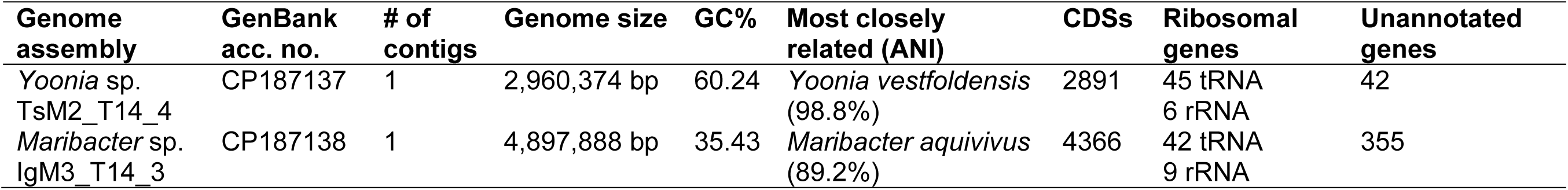
Summary of characteristics of bacterial genome assemblies of *Yoonia* sp. TsM2_T14_4 and *Maribacter* sp. IgM3_T14_3.

| Genome assembly | GenBank acc. no. | # of contigs | Genome size | GC% | Most closely related (ANI) | CDSs | Ribosomal genes | Unannotated genes |
| --- | --- | --- | --- | --- | --- | --- | --- | --- |
| *Yoonia* sp. TsM2_T14_4 | CP187137 | 1 | 2,960,374 bp | 60.24 | *Yoonia vestfoldensis* (98.8%) | 2891 | 45 tRNA<br>6 rRNA | 42 |
| *Maribacter* sp. IgM3_T14_3 | CP187138 | 1 | 4,897,888 bp | 35.43 | *Maribacter aquivivus* (89.2%) | 4366 | 42 tRNA<br>9 rRNA | 355 |

The *Yoonia* TsM2_T14_2 genome contained two PGCs harboring essential photosystem and carotenoid genes *pufABLMQ, bchBCDFHILMNXYZ, ascF, puhABCE*, and *crtABCEFI* for anoxygenic photosynthesis (Figure 2). One of the clusters (25 kb) also contained the light-sensitive regulators *ppaA* and *ppsR,* and the other cluster (15 kb) contained two *dxs* genes encoding the carotenoid precursor enzyme 1-deoxy-D-xylulose-5-phosphate synthase (DXPS).

**Figure 2.**
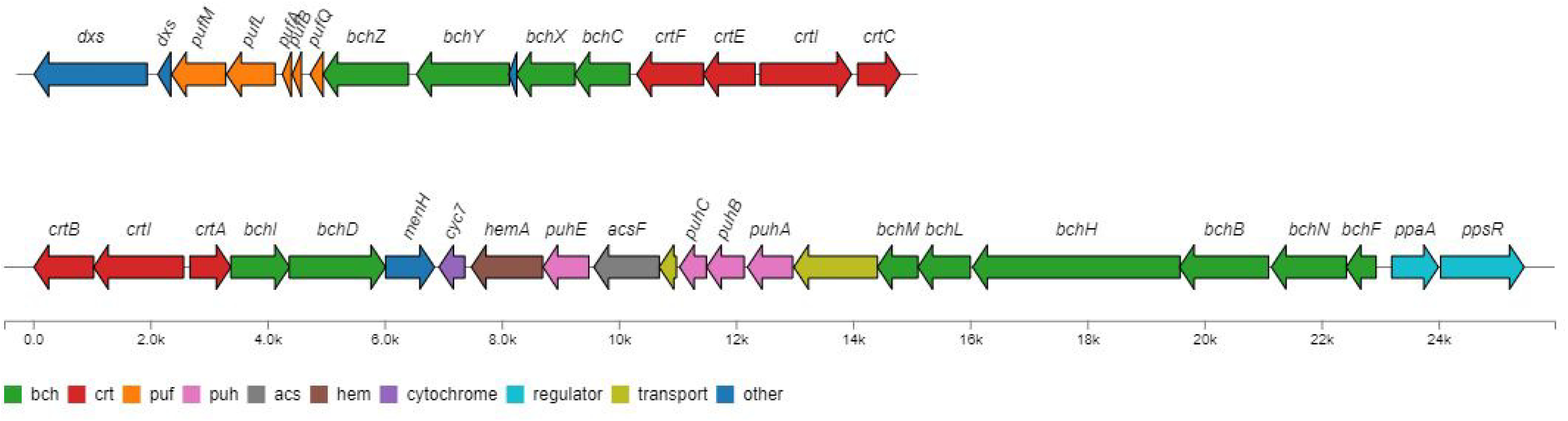
Arrangement of two putative photosynthetic gene clusters (PGCs) identified in the *Yoonia* sp. TsM2_T14_2 genome. *bch* genes (green) and *acsF* (grey) encode proteins involved in bacteriochlorophyll synthesis, *crt* genes (red) encode proteins involved in carotenoid synthesis, *puf* genes (orange) encode photosynthetic unit formation proteins, *puh* genes (pink) encode proteins involved in photosynthetic reaction center assembly, *hemA* (brown) encodes glutamyl-tRNA reductase involved in porphyrin metabolism, *cyc7* (purple) encodes cytochrome C, *ppsA* and *ppsR* (light blue) encode photosynthetically sensitive regulators, and *dxs* genes encode a synthase for a precursor of carotenoids and bacteriochlorophyll.

### Microalgal and bacterial growth in co-cultures

In the microalgal-bacterial co-cultures, microalgal cell densities started at a mean of 4.5±0.2 log10 cells ml^-1^ and increased to a mean of 4.9±0.1 log10 cells ml^-1^ for *C. weissflogii,* 6.0±0.1 log10 cells ml^-1^ for *I. galbana,* and 5.2±0.2 log10 cells ml^-1^ for *T. suecica* (Figure S1) with no significant differences between axenic cultures and co-cultures. Bacterial densities started at an average of 3.7±0.3 log10 CFU ml^-1^ for *Maribacter* sp. IgM3_T14_3, and 5.9±0.7 log10 CFU ml^-1^ for *Yoonia* sp. TsM2_T14_4. *Maribacter* sp. IgM3_T14_3 density increased the most with *C. weissflogii* to densities of 6.7±0.1 log10 CFU ml^-1^, increased to 5.9±0.2 log10 CFU ml^-1^ with *I. galbana,* and had only a slight increase to 4.1±0.3 log10 CFU ml^-1^ with *T. suecica. Yoonia* sp. TsM2_T14_4 densities also increased with all microalgal hosts, the most with *C. weissflogii* to densities of 7.3±0.1 log10 CFU ml^-1^, the least with *I. galbana* to 6.3±0.2 log10 CFU ml^-1^, and an increase to 7.0±0.2 log10 CFU ml^-1^ with *T. suecica* (Figure S2).

### Transcriptome profiles of microalgae and bacteria

Sequencing RNA from co-cultures of microalgal hosts *C. weissflogii, I. galbana,* or *T. suecica* with bacterial isolates *Maribacter* sp. IgM3_T14_3 or *Yoonia* sp. TsM2_T14_4 resulted in between 40,030,162 and 62,154,417 raw reads (PE) per sample (Table 2). After quality filtering and removal of rRNA reads, between 38,156,047 and 57,548,148 reads remained. Generally, ∼50% of reads of a relevant sample pseudoaligned to the CDSs of *C. weissflogii* and *T. suecica,* whereas around ∼15% of reads pseudoaligned to the CDSs of *I. galbana*. Bacterial reads generally were far fewer, making up 1-2% of reads in most samples except in co-cultures with *T. suecica,* where upwards of 10% of reads pseudoaligned to CDSs of *Yoonia* sp. TsM2_T14_4. In contrast, less than 1% of reads pseudoaligned to *Maribacter* sp. IgM3_T14_4, which reflected the low CFU counts of this bacterium in *T. suecica* co-cultures (Figure S2).

**Table 2.** Summary of RNA sequencing samples from algal-bacterial co-culture experiments in terms of raw reads (PE), reads after quality filtration with trimmomatic, reads that did not align to rRNA databases with SortMeRNA, and reads that pseudoaligned to algal host and bacterial CDSs, respectively, using kallisto.

| Sample | Host alga | Bacterium | Raw reads | Quality filtered reads | Non rRNA reads | Pseudoaligned to algal host CDSs | Pseudoaligned to bacterial CDSs |
| --- | --- | --- | --- | --- | --- | --- | --- |
| A1 | *Isochrysis galbana* | *Yoonia* sp. | 42,486,199 | 41,517,969 | 40,451,807 | 6,232,143 | 744,667 |
| F6 | *Isochrysis galbana* | *Yoonia* sp. | 54,547,444 | 53,019,063 | 51,710,777 | 8,009,389 | 929,109 |
| I9 | *Isochrysis galbana* | *Yoonia* sp. | 47,577,165 | 46,423,677 | 45,311,867 | 7,082,674 | 668,495 |
| X24 | *Isochrysis galbana* | *Yoonia* sp. | 59,215,772 | 57,820,715 | 56,670,087 | 9,726,220 | 958,673 |
| B2 | *Isochrysis galbana* | *Maribacter* sp. | 43,890,585 | 42,902,624 | 41,792,090 | 6,118,269 | 1,015,260 |
| C3 | *Isochrysis galbana* | *Maribacter* sp. | 52,364,932 | 51,079,743 | 49,791,277 | 7,667,074 | 1,049,398 |
| N14 | *Isochrysis galbana* | *Maribacter* sp. | 40,030,162 | 39,060,548 | 38,156,047 | 4,968,072 | 978,006 |
| R18 | *Isochrysis galbana* | *Maribacter* sp. | 46,102,099 | 45,139,287 | 44,052,005 | 5,689,066 | 1,098,000 |
| D4 | *Isochrysis galbana* | - | 45,300,181 | 44,129,896 | 42,919,128 | 6,058,148 | - |
| K11 | *Isochrysis galbana* | - | 43,046,981 | 41,915,728 | 40,782,793 | 5,911,091 | - |
| L12 | *Isochrysis galbana* | - | 53,486,243 | 52,197,292 | 50,863,948 | 6,791,448 | - |
| P16 | *Isochrysis galbana* | - | 49,442,792 | 48,121,651 | 46,959,407 | 6,310,813 | - |
| C3_2 | *Conticribra weissflogii* | *Yoonia* sp. | 53,329,066 | 52,111,257 | 50,672,567 | 26,917,211 | 747,184 |
| C7_3 | *Conticribra weissflogii* | *Yoonia* sp. | 48,531,146 | 47,426,310 | 46,045,592 | 24,716,426 | 870,328 |
| C9_5 | *Conticribra weissflogii* | *Yoonia* sp. | 43,270,929 | 42,052,547 | 40,794,109 | 21,734,653 | 630,971 |
| C34_11 | *Conticribra weissflogii* | *Yoonia* sp. | 43,120,477 | 42,047,637 | 40,585,056 | 21,359,300 | 935,332 |
| C12_6 | *Conticribra weissflogii* | *Maribacter* sp. | 53,397,676 | 52,255,366 | 50,792,084 | 26,846,979 | 1,088,298 |
| C13_7 | *Conticribra weissflogii* | *Maribacter* sp. | 40,256,395 | 39,425,264 | 38,193,376 | 20,844,956 | 574,647 |
| C26_9 | *Conticribra weissflogii* | *Maribacter* sp. | 43,983,871 | 42,943,167 | 41,691,081 | 23,591,832 | 882,749 |
| C35_12 | *Conticribra weissflogii* | *Maribacter* sp. | 46,498,588 | 45,488,052 | 44,123,800 | 23,972,230 | 900,017 |
| C2_1 | *Conticribra weissflogii* | - | 44,187,208 | 43,185,697 | 41,993,089 | 22,620,326 | - |
| C8_4 | *Conticribra weissflogii* | - | 43,468,615 | 42,509,972 | 41,297,167 | 22,756,125 | - |
| C21_8 | *Conticribra weissflogii* | - | 46,035,668 | 45,015,418 | 43,791,754 | 24,522,676 | - |
| C31_10 | *Conticribra weissflogii* | - | 47,029,762 | 45,929,506 | 44,445,627 | 24,752,467 | - |
| E5 | *Tetraselmis suecica* | *Yoonia* sp. | 50,107,303 | 48,551,072 | 46,164,798 | 21,481,182 | 3,658,607 |
| J10 | *Tetraselmis suecica* | *Yoonia* sp. | 47,438,451 | 46,013,392 | 44,151,602 | 19,889,961 | 4,316,986 |
| T20 | *Tetraselmis suecica* | *Yoonia* sp. | 54,174,843 | 52,554,494 | 50,296,623 | 23,191,989 | 4,489,643 |
| V22 | *Tetraselmis suecica* | *Yoonia* sp. | 50,893,847 | 49,370,467 | 47,426,117 | 47,426,117 | 5,180,823 |
| G7 | *Tetraselmis suecica* | *Maribacter* sp. | 48,918,860 | 47,377,333 | 45,552,745 | 21,926,329 | 39,248 |
| M13 | *Tetraselmis suecica* | *Maribacter* sp. | 45,345,422 | 43,968,396 | 41,223,706 | 18,494,905 | 68,827 |
| U21 | *Tetraselmis suecica* | *Maribacter* sp. | 53,852,172 | 52,338,173 | 49,733,219 | 20,812,784 | 97,505 |
| W23 | *Tetraselmis suecica* | *Maribacter* | 55,828,550 | 54,166,439 | 51,811,121 | 23,892,759 | 37,770 |
|  |  | sp. |  |  |  |  |  |
| H8 | *Tetraselmis suecica* | - | 62,154,417 | 60,166,239 | 57,548,148 | 26,848,903 | - |
| O15 | *Tetraselmis suecica* | - | 47,930,273 | 46,552,524 | 43,764,650 | 18,031,525 | - |
| Q17 | *Tetraselmis suecica* | - | 53,732,230 | 52,233,004 | 49,666,188 | 20,560,926 | - |
| S19 | *Tetraselmis suecica* | - | 45,551,724 | 44,249,129 | 42,534,301 | 19,544,225 | - |

Transcriptome profiles of all three host algae generally were similar regardless of the co-culture condition; axenic, or in co-culture with *Yoonia* sp. TsM2_T14_4 or *Maribacter* sp. IgM3_T14_3 (Figure 3; PERMANOVA, p-values = 0.284 (*I. galbana*), 0.217 (*T. suecica*), and 0.216 (*C. weissflogii*), perm. = 999). The transcriptomes of the two bacterial isolates, however, were significantly affected by host microalga (Figure 4) with 69.8% of the variation explained by host alga for the transcriptomes of *Yoonia* sp. TsM2_T14_4 (PERMANOVA, p-value < 0.001, perm. = 999), and 65.5% for the transcriptomes of *Maribacter* sp. IgM3_T14_3 (PERMANOVA, p-value < 0.001, perm, = 999). In total, 498 genes were differentially expressed across host algae in *Yoonia* sp. TsM2_T14_4 (adj. p-value < 0.01, |log2FC| > 1; Figure 5A). The transcriptional profile of *Yoonia* sp. with *T. suecica* as host was distinct from those when co-cultured with *I. galbana* and *C. weissflogii* as hosts. For *Maribacter* sp. IgM3_T14_3, 606 genes were differentially expressed when exposed to different host algae (adj. p-value < 0.01, |log2FC| > 1; Figure 5B), and, similarly, the transcriptional profile when with *T. suecica* was distinct from those when with *I. galbana* and *C. weissflogii*.

**Figure 3.**
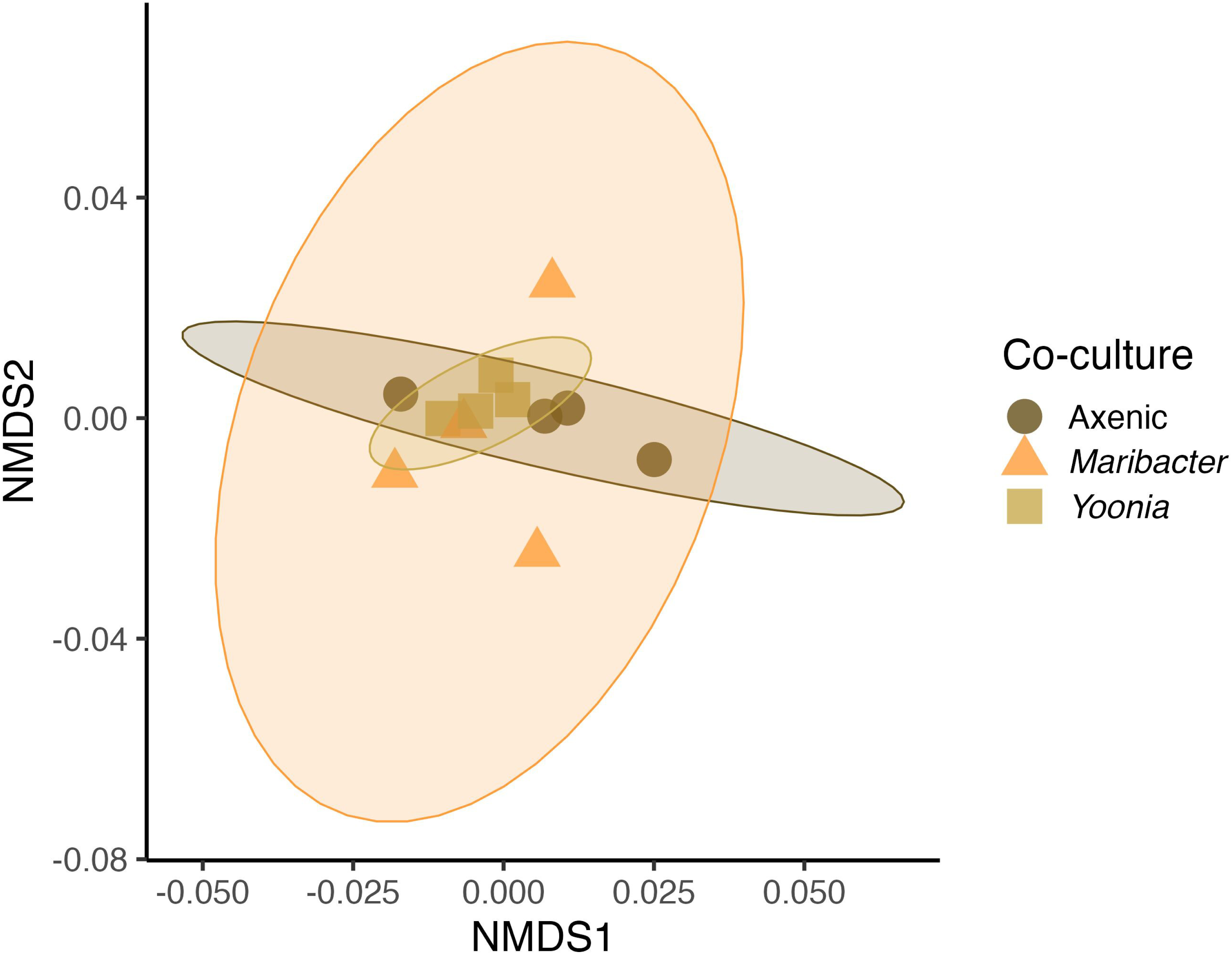
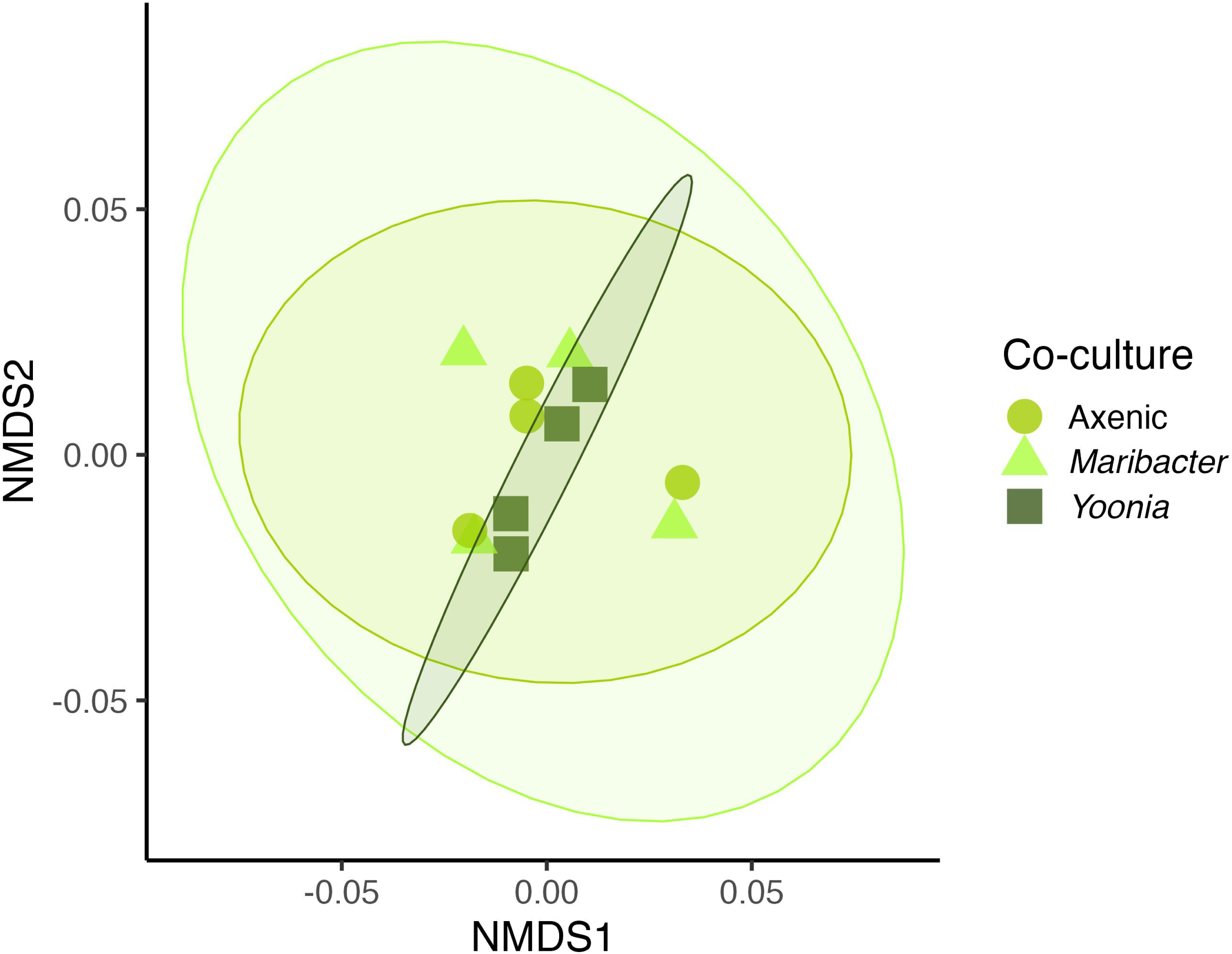
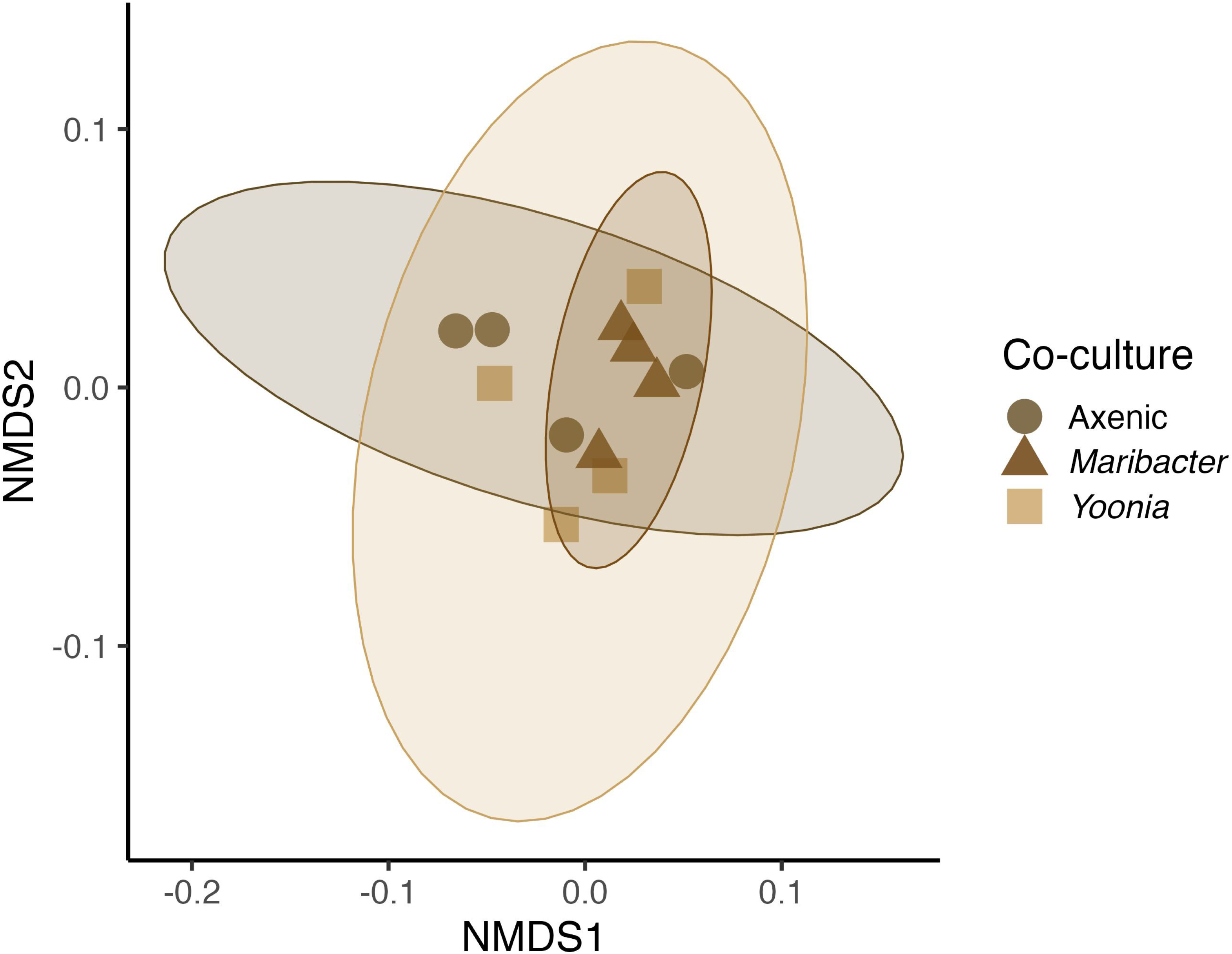
Ordination of transcriptome profiles based on Bray-Curtis dissimilarity matrix from **A***. Isochrysis galbana,* **B.** *Tetraselmis suecica,* and **C.** *Conticribra weissflogii*. Shapes and colors represent co-culture condition of a given alga, either axenic (circles), co-culture with *Maribacter* sp. IgM3_T14_3 (triangles), or co-culture with *Yoonia* sp. TsM2_T14_4m (squares). Ellipses represent 95% CIs. None of the algal transcriptomes were significantly affected by co-culture condition (PERMANOVA, p-values = 0.284 (*I. galbana*), 0.217 (*T. suecica*), and 0.216 (*C. weissflogii*), perm. = 999).

**Figure 4.**
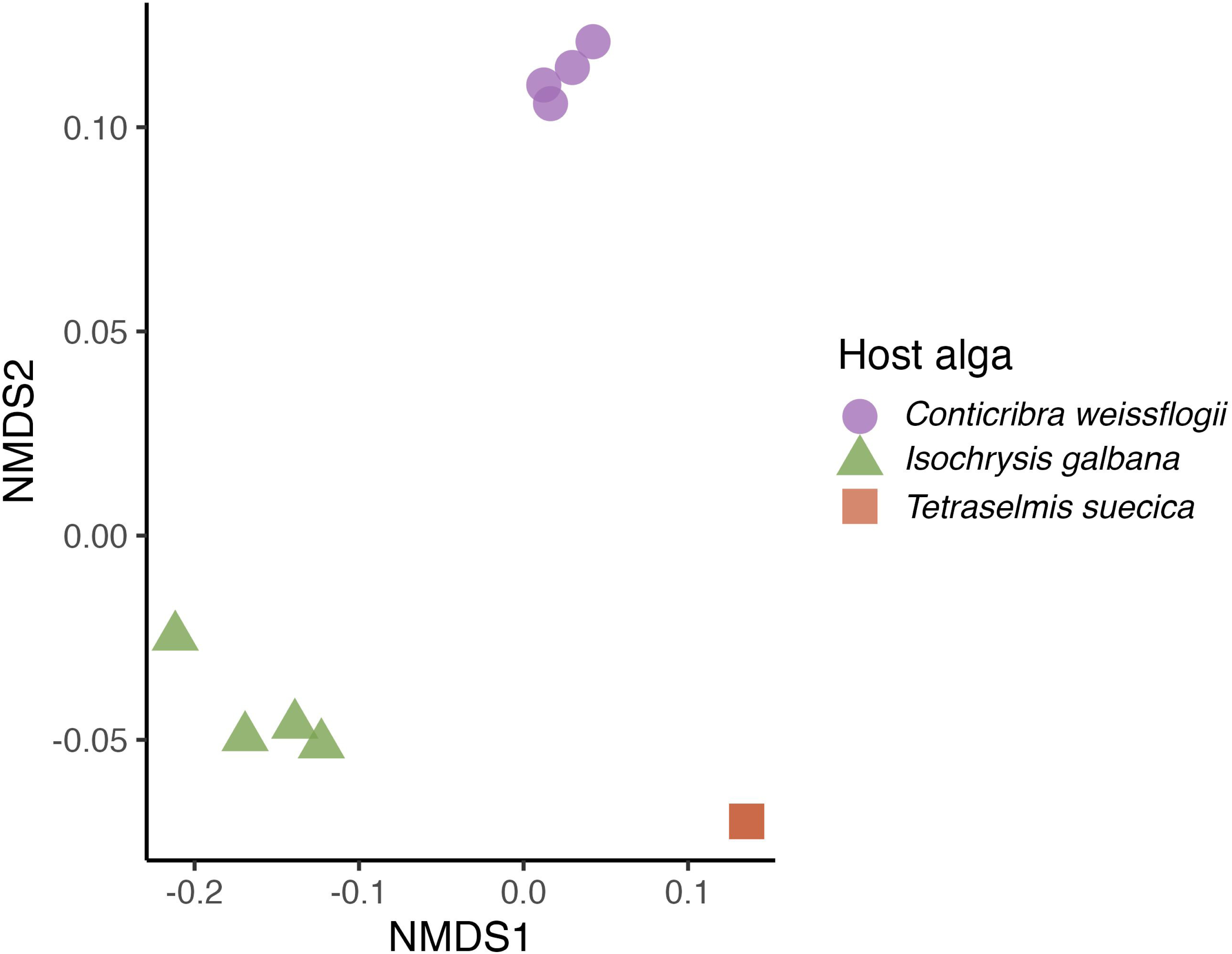
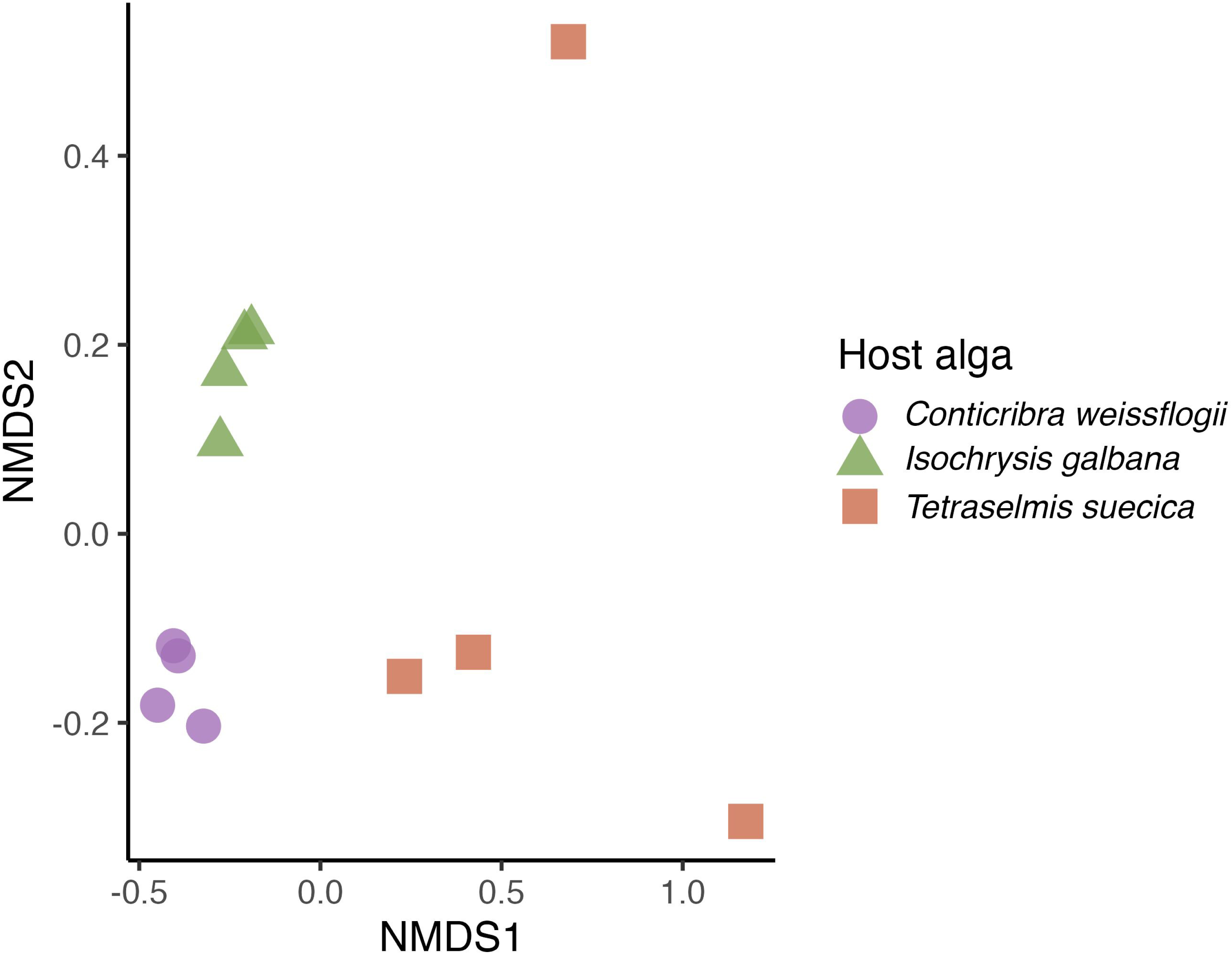
Ordination based on Bray-Curtis dissimilarity of full expression profiles of **A.** *Yoonia* sp. TsM2_T14_4 and **B.** *Maribacter* sp. IgM3_T14_3 in co-culture with three different host microalgae. Shapes and colors represent host algae *Conticribra weissflogii* (purple circles), *Isochrysis galbana* (green triangles), and *Tetraselmis suecica* (red squares). All four red squares are overlapping in 4A.

**Figure 5.**
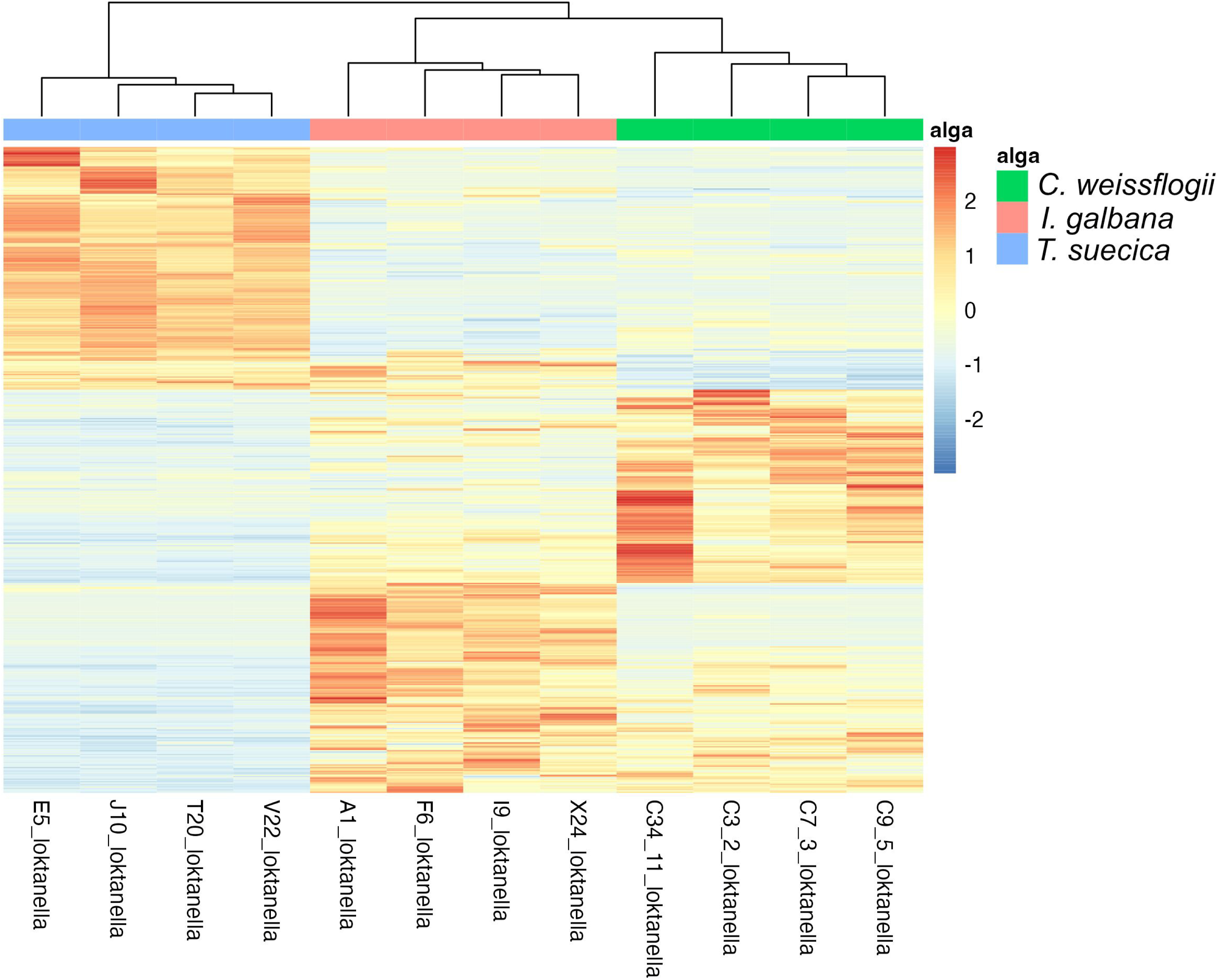
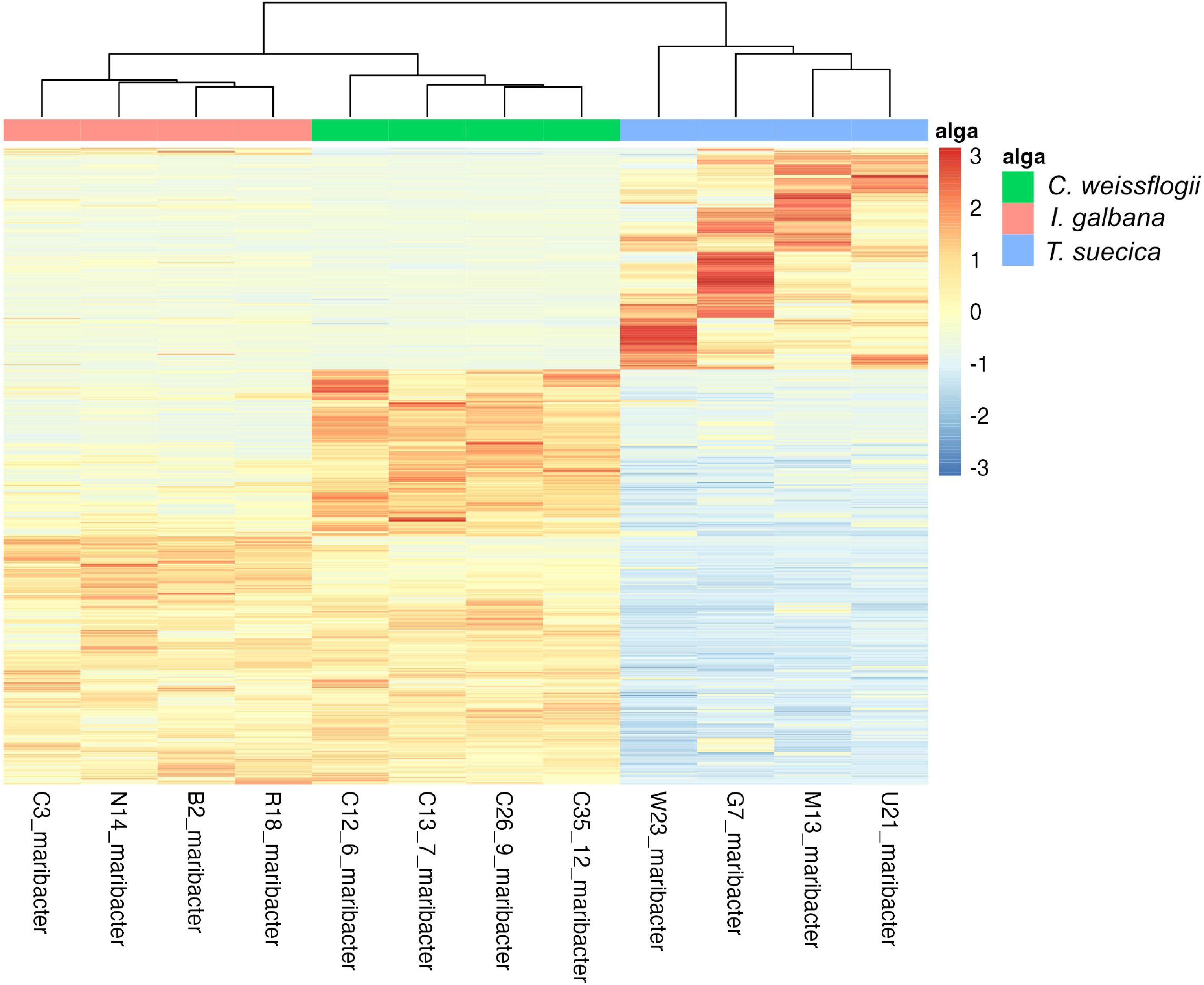
Heatmaps of differentially expressed genes (adj. p-value < 0.01 and |log2FC| > 1) across co-cultures with three host microalgae; *Conticribra weissflogii* (green bar), *Isochrysis galbana* (salmon bar), and *Tetraselmis suecica* (blue bar) in **A.** *Yoonia* sp. TsM2_T14_4 and **B***. Maribacter* sp. IgM3_T14_3. Z-scores are scaled per row.

### Interaction of *Maribacter* with host algae

Out of the differentially expressed genes in *Maribacter* (Figure 5B), transcripts putatively encoding type IX secretion system related proteins were upregulated when in co-culture with *C. weissflogii* compared to when grown with the two other microalgal hosts. Additionally, carbohydrate-related genes were significantly enriched, including alpha amylases, a glycosyl hydrolase (GH) assigned to family 16, and an alginate lyase. The GH16 amino acid sequence was further analyzed and annotated using the Conserved Unique Peptide Patterns (CUPP) webserver (98, 99). This revealed a glucan endo-1,3-beta-D-glucosidase, which can hydrolyze polysaccharides such as laminarin and curdlan. The ability of *Maribacter* IgM3_T14_3 to utilize laminarin, curdlan, or alginate as sole carbon sources was tested, and visible growth was observed with laminarin as a sole carbon source. In co-culture with *I. galbana, Maribacter* transcript counts were especially high in COG categories signal transduction mechanisms; replication, recombination, and repair; and unknown functions (Figure S3).

### Interaction of *Yoonia* with host algae

For *Yoonia* sp. TsM2_T14_4, many COG categories were highly transcribed when in co-culture with *T. suecica* compared to the two other host microalgae (Figure 5A and S4). Interestingly, among the differentially expressed genes in *Yoonia* assigned to carbohydrate transport and metabolism, a chitin deacetylase was upregulated in co-culture with *T. suecica.* The *T. suecica* draft genome does harbor three putative chitin synthase genes, of which all three were expressed in the co-culture experiment. However, chitin utilization assays showed no degradation of colloidal or crystalline chitin by *Yoonia* sp. TsM2_T14_2.

When in co-culture with *C. weissflogii,* highly transcribed genes of *Yoonia* sp. TsM2_T14_2 belonged to COG categories nucleotide transport and metabolism; lipid transport and metabolism; and unknown functions. Other enriched transcripts in co-culture with *C. weissflogii* were related to genes involved in cobalamin synthesis and metabolism. This finding generated the hypothesis that *Yoonia* sp. TsM2_T14_2 could be supplying cobalamin to *C. weissflogii* when in co-culture. However, the addition of *Yoonia sp.* TsM2_T14_2 to a *C. weissflogii* culture without cobalamin did not result in higher algal growth rates or algal cell density compared to a culture with no bacterial symbiont. In fact, the presence of *Yoonia* sp. TsM2_T14_2 in a cobalamin-depleted co-culture caused a decrease in cell density of *C. weissflogii* in late growth stages (day 20, Figure S5).

Finally, in co-culture with *I. galbana*, COG categories highly transcribed were transcription; cell cycle control, cell division, chromosome partitioning; and carbohydrate transport and metabolism. Additionally, genes related to photosynthesis (light-harvesting antenna proteins, photosynthetic reaction center subunits, and chlorophyll synthesis proteins) and phages (phage tail proteins, phage proteins, large subunit phage terminase) were upregulated in *Yoonia* sp. TsM2_T14_2 when in co-culture with *I. galbana*.

## Discussion

In the present study, we compared expression profiles of microalgal hosts and of two associated bacterial isolates to investigate if the phycosphere bacteria confer a benefit to their microalgal hosts and/or vice versa. When comparing expression profiles of *I. galbana, T. suecica,* and *C. weissflogii* as axenic culture to those in co-culture with *Maribacter* sp. IgM3_T14_3 or *Yoonia* sp. TsM2_T14_4, none of the algal expression profiles showed significant differences. This is in contrast to previous studies, where microalgal host *Chlorella* spp. C596 exhibited specific transcriptional responses to different microbiomes (31) as well as studies of the metabolome of *Thalassiosira pseudonana* in response to individual bacterial isolates (100). Additionally, these results suggest (under the conditions tested here) that the algal hosts are not actively interacting with these bacterial partners. While the volume of studies systematically assessing the effect of particular bacteria on microalgal hosts remains small, one study observed a majority of neutral or positive effects on algal growth or carrying capacity by 368 phycosphere isolates (101). In contrast, another study found mostly neutral or antagonistic effects of bacterial isolates on a diatom host (13). In the present study, we show a neutral effect of common phycosphere bacteria on microalgal hosts not only in terms of effects on algal growth but also on algal gene expression. This observation suggests that even though some bacteria are part of the core microbiome of certain microalgae, they might not confer an effect on the host gene expression and growth under all conditions. It should be noted, however, that these bacteria may confer effects on the microalgal hosts under other circumstances than the experimental conditions tested here, for instance, under xenic conditions where the bacteria may interact with other members of the phycosphere microbiome. Given these results and those of Baker et al. (2023) and Stock et al. (2019), perhaps neutral effects of phycosphere bacteria on microalgal hosts are more common than often assumed, and many interactions in the phycosphere are commensal (bacteria benefitting) rather than mutually beneficial or antagonistic.

The expression profiles of *Maribacter* sp. IgM3_T14_3 and *Yoonia* sp. TsM2_T14_4 did differ significantly when co-cultured with different microalgal hosts. It should be noted that these differences could be impacted by differences in growth of the bacteria in the co-cultures, in particular the fact that *Maribacter* did not reach high cell densities in co-culture with *T. suecica* compared to the other microalgal hosts. Keeping this in mind, the differential responses observed here correspond to previous findings of expression profiles of another Rhodobacteraceae representative, *Ruegeria pomeroyi,* a Flavobacteriaceae representative, *Polaribacter dokdonensis,* and a *Stenotrophomonas* sp., that responded differently to the diatom host *Thalassiosira pseudonana* and the chlorophyte host *Micromonas commoda* (41, 42).

In the transcriptome of *Maribacter* sp. IgM3_T14_3 in co-culture with *C. weissflogii,* several genes putatively involved in interactions with the host were upregulated, including genes involved in carbohydrate transport and metabolism and genes related to the type IX secretion system (T9SS). Interestingly, this has also been observed in another Bacteroidota, a *Dyadobacter* sp. with microalgal hosts *Micrasterias radians* and *Scenedesmus quadricauda* (11, 44). While the T9SS can be involved in a variety of processes including virulence and motility, it can also be involved in the secretion of carbohydrate-degrading enzymes such as agarases, cellulases, chitinases, and xylanases (102). Here, we show that *Maribacter* sp. IgM3_T14_3 can utilize laminarin as a sole carbon source and that it highly expressed a GH16 glucan endo-1,3-beta-D-glucosidase in co-culture with *C. weissflogii*. Hence, we hypothesize that this enzyme is secreted through the T9SS of *Maribacter* sp. IgM3_T14_3 to hydrolyze laminarin into smaller saccharides before transport into the cell. Since *C. weissflogii* does produce laminarin under conditions similar to those tested in the present study (103), it is likely that this could be the case here as well and that *Maribacter* sp. IgM3_T14_3 can benefit from laminarin produced by *C. weissflogii* as a carbon source.

In the *Yoonia* sp. TsM2_T14_4 expression profiles, *Yoonia* overexpresses a gene involved in chitin degradation, a chitin deacetylase-encoding gene (104) when in co-culture with *T. suecica*. While chitin is a known part of cell walls in fungi and diatoms (Durkin et al., 2009; Shao et al., 2019), it has not thus far been reported as a major component in chlorophytes. Here, however, three putative chitin synthases in the *T. suecica* genome were expressed under all experimental conditions, suggesting that *T. suecica* might be producing chitin constitutively. Hence, a core phycosphere microbiome member such as *Yoonia* sp. TsM2_T14_4 could benefit from degrading chitin produced by *T. suecica*. However, we were unable to demonstrate chitin (crystalline nor colloidal) degradation capabilities of *Yoonia* sp. TsM2_T14_4. It is possible that the laboratory conditions tested here were not sufficient to detect chitin degradation, or that the overexpressed enzyme is involved in the degrading polysaccharides other than chitin. Furthermore, chitin differs largely in structure (chitin type, degree of polymerization and acetylation) (105, 106), and chitin from *T. suecica* may be of a different structure than the crab or shrimp chitin utilized in the chitin degradation assay. Further work will be required to determine the exact function of this overexpressed gene and if it is important in the interaction of *Yoonia* sp. TsM2_T14_4 with *T. suecica* or other algal hosts.

In co-culture with *I. galbana*, *Yoonia* sp. TsM2_T14_4 upregulated genes related to anoxygenic photosynthesis, a metabolic capability which has previously been found in related roseobacters such as *Yoonia vestfoldensis* (107, 108). While generally underexplored, it has been proposed that this process serves to support the bacterium’s energy needs during periods of nutrient scarcity (109, 110). It is possible that *Yoonia* sp. TsM2_T14_4 is experiencing starvation in co-culture with *I. galbana* due to its poor degradation capabilities of HMW carbohydrates (38, 39) and hence resorts to anoxygenic photosynthesis. This could also explain why it was outcompeted by other bacterial taxa in the microbiomes of *I. galbana* previously characterized (24) due to the high energetic burden of synthesis of photosynthetic units (111). However, further experimental work is needed to clarify whether this is the case.

*Yoonia* sp. TsM2_T14_4 grew to high cell densities in co-culture with all three microalgal hosts, whereas *Maribacter* sp. IgM3_T14_3 grew to high densities with *I. galbana* and *C. weisssflogii* as hosts, but only reached low cell densities in co-culture with *T. suecica*. In our previous study (24), *Maribacter* sp. was a core part of the *I. galbana* microbiome, but not of the other two host microalgae, conflicting with the growth patterns observed here. Similarly, *Yoonia* sp. was a core part of *T. suecica* and *C. weissflogii* microbiomes but reached high cell densities with all algal hosts in the current study. These growth patterns point to other interactions impacting the growth and prevalence of *Maribacter* sp. and *Yoonia* sp. in *I. galbana, T. suecica,* and *C. weissflogii* microbiomes. In more complex systems, it is possible that higher-order interactions between more than two partners govern growth patterns and interactions (112, 113), highlighting this limitation of the simpler co-culture systems studied here.

In conclusion, the interactions between phycosphere bacteria *Maribacter* sp. IgM3_T14_3 or *Yoonia* sp. TsM2_T14_4 and microalgal hosts *C. weissflogii, I. galbana,* or *T. suecica* are complex and not fully understood. It appears that the interactions are mostly commensal with the microalgal hosts being largely unaffected by the presence of *Maribacter* or *Yoonia* when compared to an axenic state. The dual transcriptomics approach presented here can aid in revealing phycosphere interactions and generating hypotheses for experimental follow-up as exemplified here. Carbohydrate exudates from the microalgal hosts seem to shape the transcriptional response of *Maribacter* and *Yoonia,* e.g. laminarin produced by *C. weissflogii* being degraded by *Maribacter*, and *Yoonia* presumably lacking capabilities in degrading carbohydrates from *I. galbana* and utilizing anoxygenic photosynthesis as an alternative or complementary energy pathway. However, a large range of responses to other compounds released by the microalgae also shape the expression profiles of *Maribacter* and *Yoonia* and are not well understood at this time. As model phycosphere bacteria, *Maribacter* for Rhodobacteraceae and *Yoonia* for Flavobacteriaceae, these two phycosphere isolates show a varied transcriptional response to the three microalgal hosts, demonstrating different, but well-adapted strategies to life in the phycosphere.

## Acknowledgements

The authors would like to thank Peter Bing Svendsen for assisting with ONT sequencing of the genomes of the two bacterial isolates, Mikael Lenz Strube for helpful discussions on bioinformatics pipeline, and Jesper Holck for discussions about carbohydrate degradation and providing polysaccharides for degradation tests. This project was funded by the Novo Nordisk Foundation (NNF20OC0064249) that also provided equipment funding (NNF19OC0055625).

